# Visual field asymmetries develop throughout adolescence

**DOI:** 10.1101/2023.03.04.531124

**Authors:** Marisa Carrasco, Caroline Myers, Mariel Roberts

**Affiliations:** Dept. of Psychology, New York University, New York, NY 10003, USA; Center for Neural Science, New York University, New York, NY 10003, USA; Dept. of Psychological & Brain Sciences, Johns Hopkins University, Baltimore, MD 21218, USA; Dept. of Psychology, Barnard College, New York, NY 10027

## Abstract

For human adults, visual perception varies with polar angle at isoeccentric locations from the center of gaze. The same visual information yields better performance along the horizontal than vertical meridian (horizontal vertical anisotropy, HVA) and along the lower than upper vertical meridian (vertical meridian asymmetry, VMA). For children, performance is better along the horizontal than vertical meridian (HVA) but does not differ between the lower and upper vertical meridian. Here, we investigated whether the extent of the HVA varies and whether the VMA emerges and develops during adolescence, or whether the VMA only emerges in adulthood. We found that both the HVA and VMA develop gradually throughout adolescence and become as pronounced as those of adults only in late adolescence.

## INTRODUCTION

Visual perception in human adults varies throughout the visual field. It decreases across eccentricity –distance from the center of gaze– and changes around isoeccentric locations –with polar angle at a constant distance from the center of gaze. The same stimuli yield better performance along the horizontal than vertical meridian (HVA) and along the lower than upper vertical meridian (VMA). In adults, these perceptual polar angle asymmetries, and their retinal and cortical correlates, have been well characterized (review, Himmelberg, Winawer & Carrasco, 2023).

For adults, polar angle asymmetries are pervasive; they are present for basic dimensions, such as contrast sensitivity (e.g., Carrasco, Talgar & Cameron, 2001; Cameron, Tai & Carrasco, 2002; Himmelberg, Winawer & Carrasco, 2020; Jigo & Carrasco, 2020; Pointer & Hess, 1989; Rijsdijk, Kroon & Van der Wildt, 1980; Rovamo & Virsu, 1979) and acuity (Barbot, Xue & Carrasco, 2021; Montaser-Koushari & Carrasco, 2009; Kwak, Hanning & Carrasco, 2023).

These asymmetries become more pronounced as stimulus spatial frequency and eccentricity increase and when the target appears amidst distractors (Carrasco et al., 2001; Purokayastha, Roberts & Carrasco, 2021). They are present across stimulus properties, e.g., luminance, contrast, orientation, size, spatial frequency (Carrasco et al., 2001; Himmelberg et al., 2020; Kwak et al., 2024), when stimuli are viewed monocularly or binocularly (Carrasco et al., 2001; Barbot et al., 2021), regardless of whether stimuli are masked or not (Carrasco et al., 2001; Carrasco, Williams & Yeshurun, 2002), stimulus orientation and tilt angle (Baldwin, Meese & Baker, 2012; Barbot et al., 2021; Purokayastha et al., 2021) and head orientation. Both HVA and VMA are retinotopic: when observers rotate their head they shift with the stimulus retinal location, not with its location in space (Corbett & Carrasco, 2011). Interestingly, their extent increases with eccentricity from center of gaze (Baldwin et al., 2012; Carrasco et al., 2001; Fuller, Rodriguez & Carrasco, 2008; Jigo et al., 2023) and with spatial frequency (Barbot et al., 2021; Carrasco et al., 2001; Himmelberg et al., 2020). Furthermore, these asymmetries are strongest at the cardinal meridians and diminish gradually with angular distance from them, so that they are no longer present at the intercardinal meridians (Abrams, Nizam & Carrasco, 2012; Baldwin et al., 2012; Barbot et al., 2021; Carrasco et al., 2001).

These asymmetries are also present for mid- and higher-level vision, e.g. texture segmentation (Talgar & Carrasco, 2002), illusory contours (Rubin et al., 1996), letter recognition (Mackeben, 1999), perceived size (Schwarzkopf, 2019), face perception (Roux-Sibilon et al., 2023), and numerosity processing (Chakravarthi, Papadai & Krajnik, 2022), as well as in visual short-term memory (Montaser-Koushari & Carrasco, 2009).

Optical quality and retinal factors (e.g., cone density and midget retinal ganglion cell density) account for a small percentage of these perceptual asymmetries (Kupers, Carrasco, & Winawer, 2019; Kupers et al., 2022). These asymmetries are better accounted for by the distribution of cortical surface area in primary visual cortex (V1), which is larger for the horizontal than vertical meridian, and for the lower than upper vertical meridian (Silva et al., 2018; Benson et al., 2021; Himmelberg et al., 2022; Himmelberg et al., 2021, 2023). Moreover, individual differences in contrast sensitivity at the cardinal meridians correlate with the V1 surface representing these meridians (Himmelberg, Winawer & Carrasco, 2022).

For children (5-12 years), visual performance yields an HVA, albeit less pronounced than that for adults, but not a VMA (Carrasco et al., 2022). Correspondingly, children’s cortical surface is larger for the horizontal than vertical meridian but is similar for the upper and lower vertical meridian (Himmelberg et al., 2023). In addition to these ontogenetic differences, phylogenetic differences exist: Whereas human adults show the typical VMA in a motion discrimination task, non-human adult primates show an inverted VMA; discrimination is better for stimuli at the upper than the lower vertical meridian (Tünçok, Kurpes & Carrasco, 2025).

Here, we investigated whether the extent of the HVA varies and the VMA emerges and fully develops during adolescence, or whether the VMA only emerges in adulthood. Adolescents aged 13–17 years (n=155) and adults (n=112) performed an orientation discrimination task. (Data for adults have been reported in a recent study (Carrasco et al., 2022). Gabor patch stimuli appeared briefly (∼100 ms) at four isoeccentric locations along the cardinals. Participants indicated whether the target was tilted right or left from vertical (**Figure 1**). Performance (d’) was calculated at each location according to signal detection theory.

**Figure 1.**
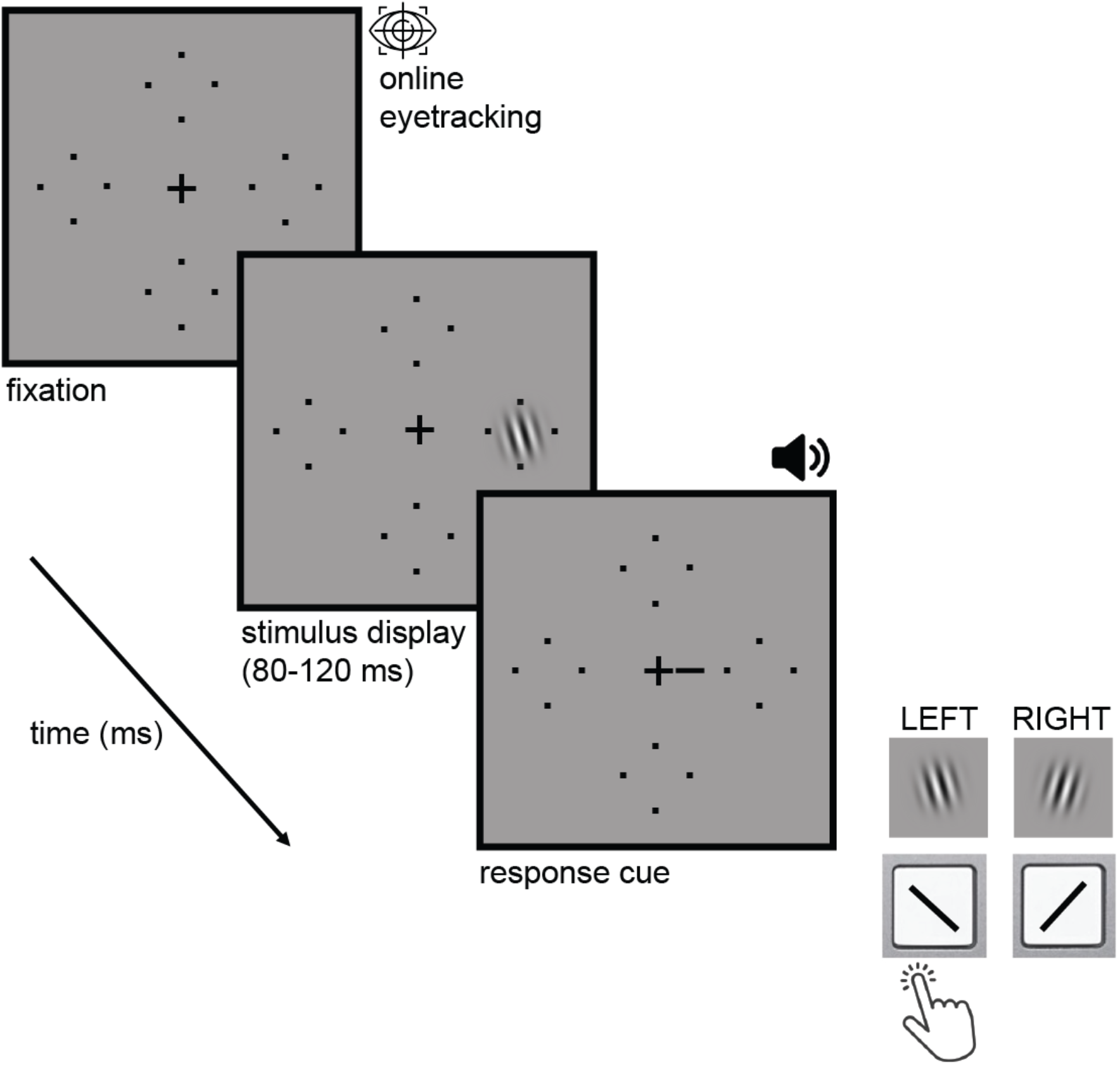
Experimental trial: orientation discrimination task. Observers fixated on the plus sign in the display center. Stimuli were briefly presented at four isoeccentric locations. A response cue indicated the target location, and observers reported its orientation (left vs. right) with a key press. Observers’ eyes were tracked to ensure stable fixation. A feedback tone indicated a correct or incorrect response.

## RESULTS

Adolescents’ discriminability (d’) revealed both an HVA and VMA, as did adults’ discriminability (**Figure 2A,C**). The d’ HVA was more pronounced for adults than adolescents. A 2×2 (age X location) analysis of variance (ANOVA) revealed an interaction (F(1,265) =28.26, p<0.0001). The superior performance on the horizontal than vertical meridian was more pronounced for adults (F(1,111) =142.89, p<0.0001) than adolescents (F(1,154) =48.48, p<0.0001).

**Figure 2.**
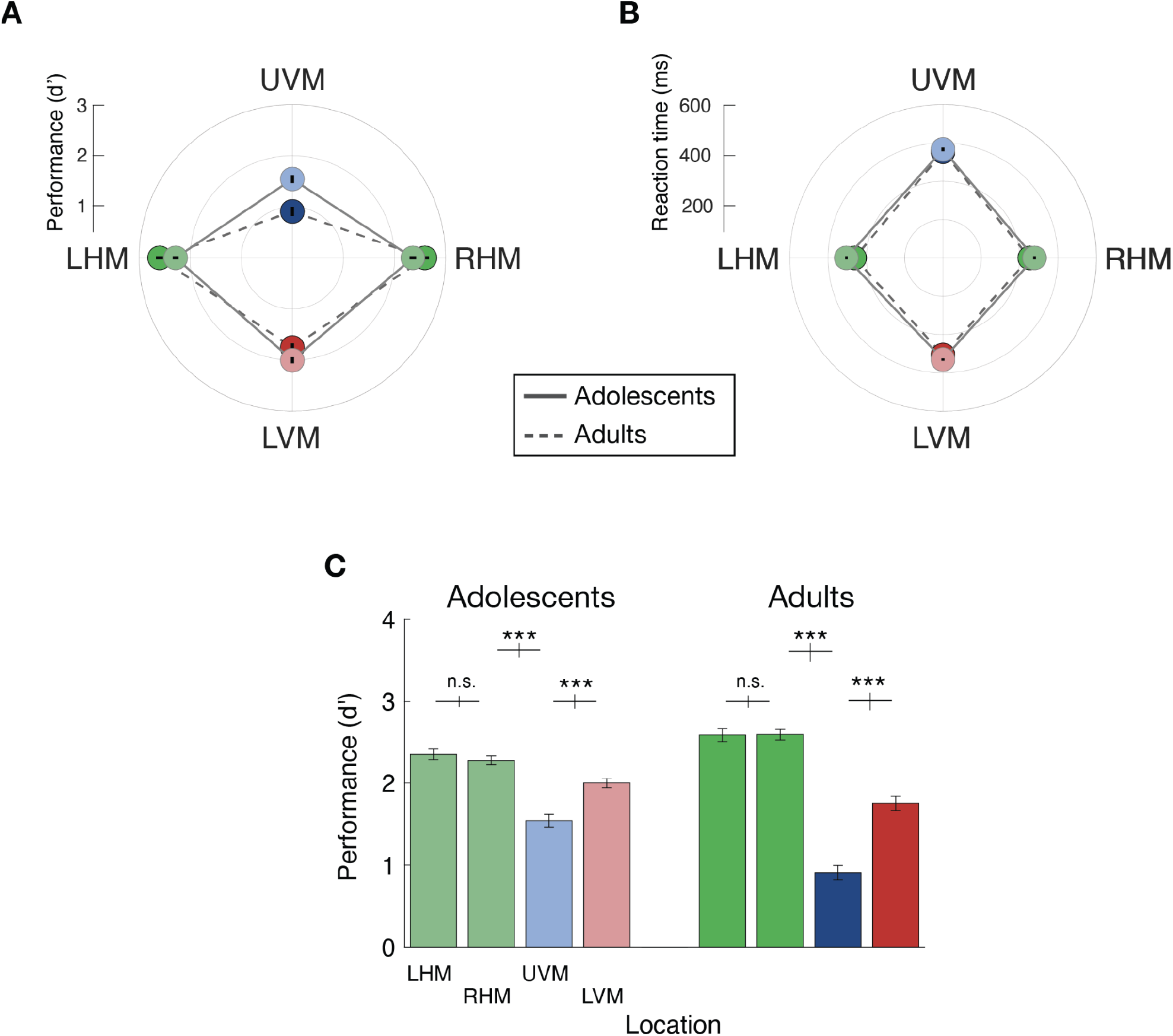
Polar angle asymmetries differ between adolescents and adults. **(A)** Adolescents’ (n=155) and adults’ (n=112) polar angle asymmetries in visual performance (d’). The target was presented at one of four isoeccentric cardinal locations on each trial. The center point represents chance performance and each of the four points represents discriminability (d’) at those locations, with the black line inside representing ±1 SEM. For adults (dashed lines), performance shows the typical asymmetries: (1) horizontal–vertical anisotropy (HVA): better performance along the horizontal than vertical meridian; and (2) vertical–meridian asymmetry (VMA): better performance at the location directly below (LVM) than above (UVM) fixation. For adolescents (solid line), both asymmetries are present, but less pronounced. **(B)** Adolescents’ and adults’ polar angle asymmetries in response time (ms). HVA and VMA are present in both groups. Neither asymmetry interacted with age. **(C)** Orientation discriminability (d’) at each location: left-horizontal meridian (LHM), right-horizontal meridian (RHM), upper vertical meridian (UVM) and lower vertical meridian LVM) for adolescents and adults. In **A** and **B**, error bars = ±1 within-subject SEM. In **C**, error bars between locations = standard error of the difference *=p<.05, **=p<0.01, ***=p<0.001.

The d’ VMA also differed between age groups (**Figure 2A,C**): a 2×2 (age x location) ANOVA revealed an interaction (F(1,265) =6.99, p<0.01). The superior performance on the lower than upper vertical meridian was more pronounced for adults (F(1,111) =62.9, p<0.0001) than adolescents (F(1,154) =29.16, p<0.0001).

The HVA and VMA were paralleled in response times (**Figure 2B**). Both groups had both asymmetries: HVA in adults (F(1,111) =51.983, p<.0001) and adolescents (F(1,154) =48.847, p<.0001); VMA in adults (F(1,111) =17.79, p<.0001) and adolescents (F(1,154) =11.499, p<.001). There was no age x location interaction for either the HVA (F(1,265) =1.61, p>0.1) or the VMA (F(1,265) =.55, p>0.1). Thus, there were no speed-accuracy trade-offs in either adolescents or adults and no difference in RT between groups.

To assess the developmental trajectory of the HVA and emergence of the VMA, we divided the adolescent participants into 5 groups according to their age and analyzed whether the extent of these asymmetries differed for these subgroups (**Figure 3A**): An HVA was present for all ages after 14 years and became more pronounced with age. A 5×2 (age X location) analysis of variance (ANOVA) revealed an effect of location (F(1,150) =41.62, p<0.0001) and an interaction with age (F(4,150) =3.409, p<0.05). The VMA also marginally interacted with age (F(4,150) =2.383, p=0.054).

**Figure 3.**
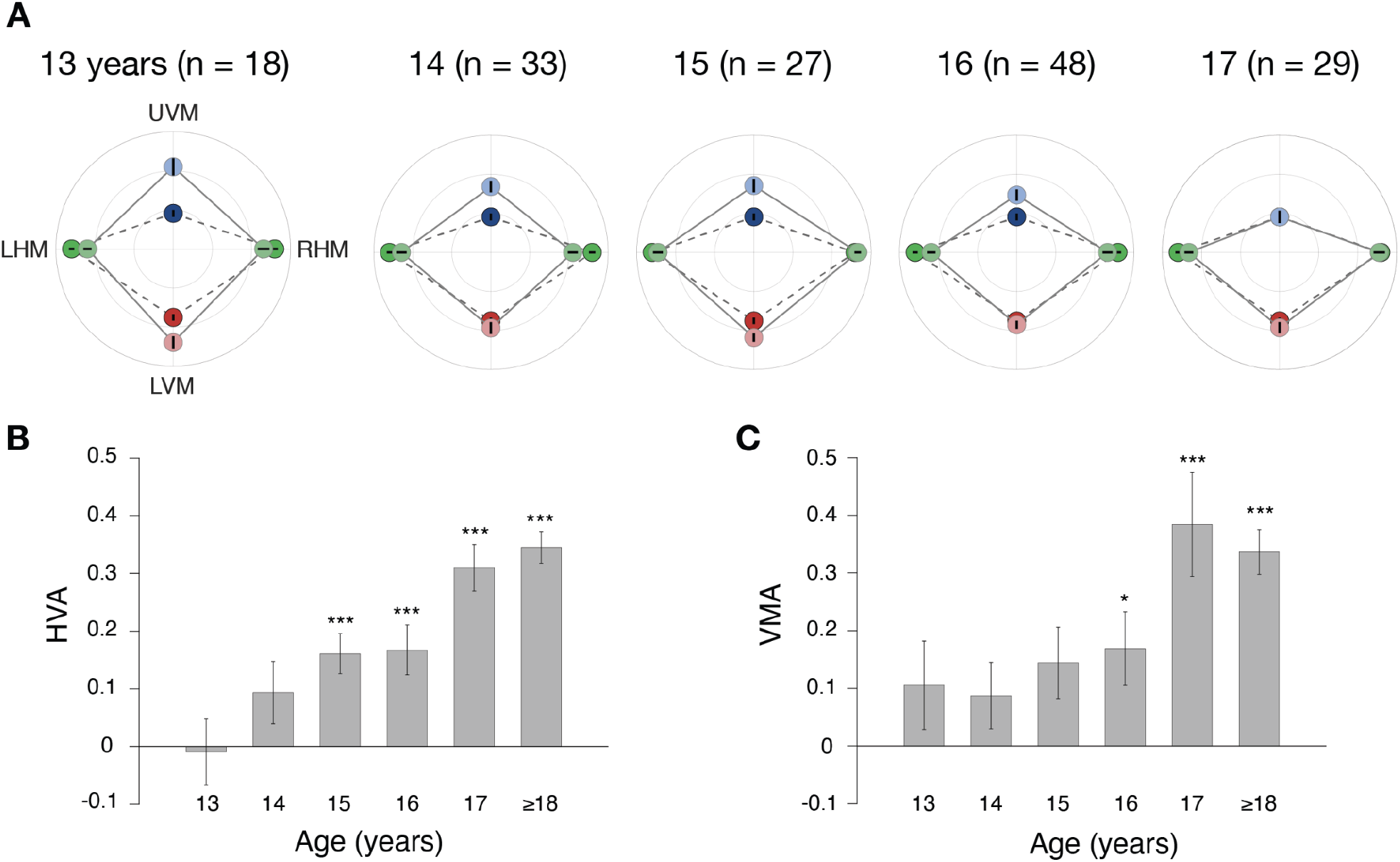
Polar angle asymmetries per adolescents’ age groups. (**A**) Polar angle asymmetries per age group (same format as in (**2A**)). Data for adults (dashed lines) have been published in Carrasco et al. (2022). (**B, C**) Magnitudes of the (B) HVA ratios [(Horizontal–Vertical)/(Horizontal+Vertical)] and (C) VMA ratios [(Lower Vertical–Upper Vertical)/(Lower Vertical+Upper Vertical)] for each group of adolescents and adults (asterisks reflect within-adolescent Bonferroni-corrected significance). Data for adults (≥18) have been published in Carrasco et al. (2022). Error bars = ±1 within-subject SEM. *=p<.05, **=p<0.01, ***=p<0.001.

The magnitude of the HVA [(Horizontal–Vertical)/(Horizontal+Vertical); **Figure 3B**] differed for adolescents and adults (F(1,265) =29.02, p<.0001); the asymmetry was less pronounced for adolescents (t(154)= 7.05, p<.0001) than adults (t(111) =12.586, p<.0001). Moreover, HVA ratios differed among adolescent age groups (F(4,150) =4.72, p=.001), becoming significant by age 15 (t(26) =4.648, p bonf <.001), and increasing with age.

Likewise, the magnitude of the VMA [(Lower Vertical–Upper Vertical)/(Lower Vertical+Upper Vertical; **Figure 3C**] differed for adolescents and adults (F(1,265) =9.51, p<.01); the asymmetry was less pronounced for adolescents (t(154) =5.519, p<.0001) than adults (t(111) =8.66, p<.0001). Moreover, the VMA differed among adolescent age groups (F(4,150) =2.57, p=.041), becoming significant by age 16 (t(47) =2.64, p bonf <.05), and increasing with age.

We also plotted individual values for each subject in each group level for the HVA and VMA. The HVA becomes significant at age 15 and is more consistent by age 17 (**Figure 4A**). The VMA becomes significant at age 16, and is more pronounced and consistent by age 17 and into adulthood (**Figure 4B**).

**Figure 4.**
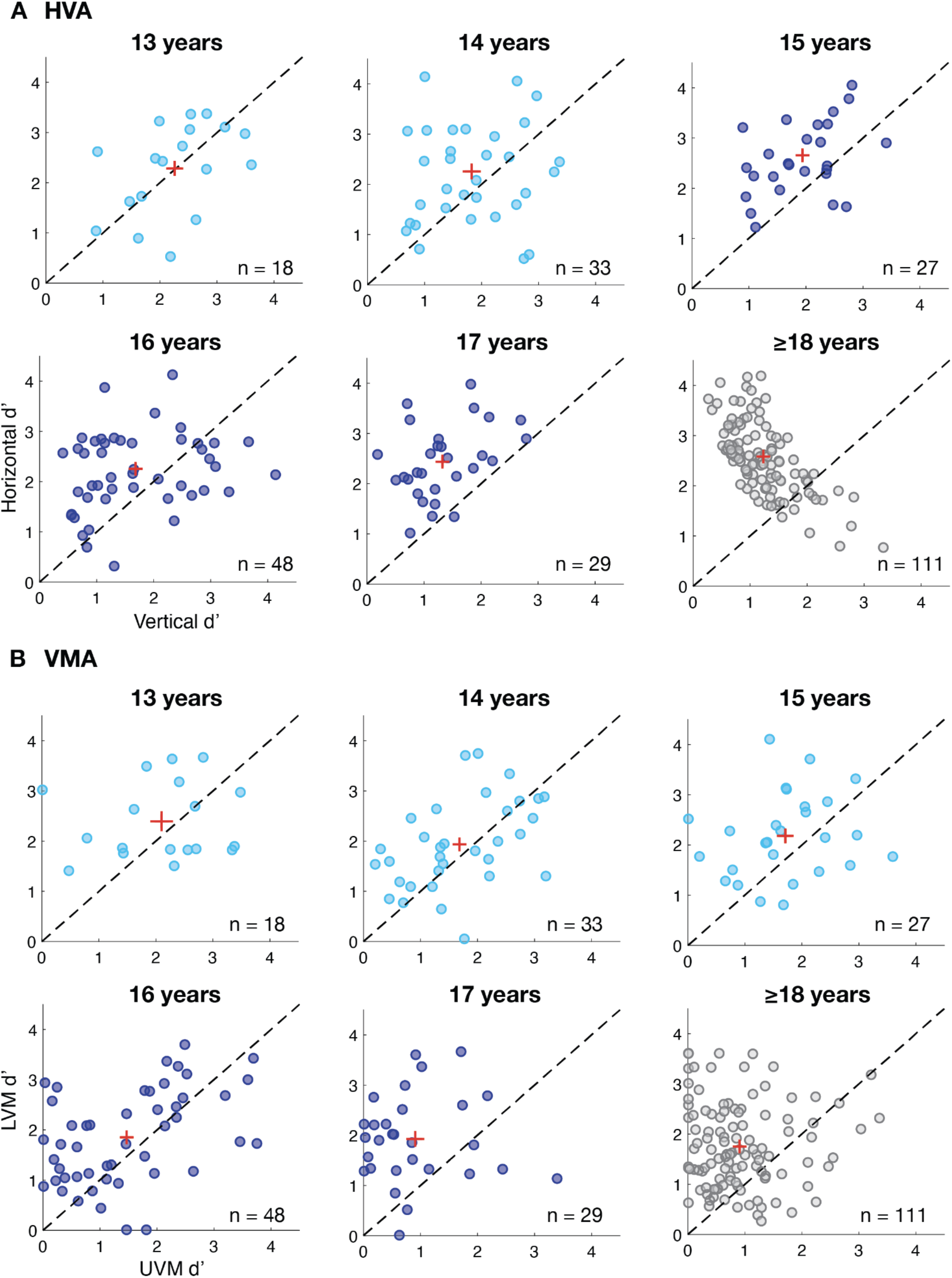
Individual asymmetry ratios for each age group. **(A)** The HVA for each adolescent subgroup and for adults. For each age group, data above the diagonal indicate better performance at the horizontal than vertical meridian and vice versa. **(B)** The VMA for each adolescent subgroup and for adults. For each group, data above the diagonal indicate better performance at the lower than upper vertical meridian and vice versa. Lighter blue panels indicate age groups that did not show significant asymmetries; darker blue panels indicate age-groups that showed significant asymmetries. The center of the red cross indicates the mean value for each group and their line lengths represent ±1 SEM.

To evaluate whether height contributes uniquely to the magnitude of the HVA and VMA independently from age, we regressed out age from these measures and then correlated the residuals with height (e.g., Himmelberg et al., 2022; Lee & Carrasco, 2025). These analyses included the 347 observers (91% of the total number of adolescents, children, and adults) for whom height was available. We found no significant correlation for HVA (Spearman’s r= .039, p>.1; **Figure 5A**). In contrast, height correlated with VMA residuals (Spearman’s r= .188, p<.001; **Figure 5B**), demonstrating a significant association between observers’ height and VMA magnitude when age-related variance is removed.

**Figure 5.**
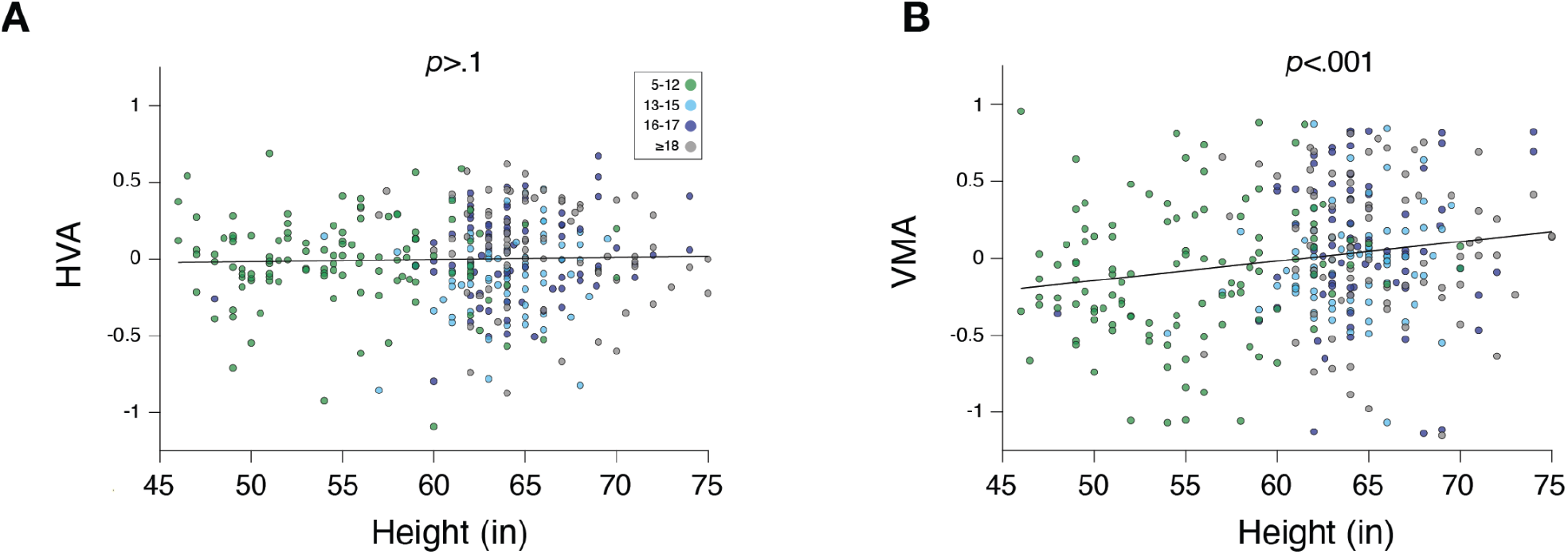
Correlations between individuals’ asymmetry ratios and height. **(A)** Correlation between (age-residualized) HVA ratios and height for 347 observers, Spearman’s r=.039, p>.1. **(B)** Correlation between (age-residualized) VMA ratios and height for 347 observers, Spearman’s r=.189, p<.001. HVA and VMA values for observers aged 5-12 and ≥18 ys old have been reported in Carrasco et al. (2022).

## DISCUSSION

In this study we found that both the HVA and the VMA develop gradually and independently. The HVA, which is already present for children behaviorally (Carrasco et al., 2022) and cortically (Himmelberg et al., 2023), becomes stronger with age and seems to be well established by 15 years old. The VMA, which is not present in children (Carrasco et al., 2022; Himmelberg et al., 2023), only emerges in adolescence and seems to be well established by 17 years of age, towards its end.

These systematic differences in performance around the visual field throughout adolescence and between adolescents and adults provide further evidence that some basic aspects of visual perception and the brain continue to develop until the start of adulthood (Benedek et al., 2010; Dekker et al., 2020; Linka, Karimpur & de Haas, 2025; Toga, Thompson & Sowel, 2006).

For instance, from ages 4 to 18, the contrast sensitivity function (CSF) improves at an exponentially decaying rate by approximately 0.3 log_10_ units, corresponding to a doubling of contrast sensitivity. Although 90% of this change appears to be reached by age 12 (Dekker et al., 2020), our findings show that this improvement is not homogeneous around the visual field. Were the enhancement uniform, the HVA and VMA would be equally present across ages, rather than vary developmentally.

A recent study has also reported a protracted increase in the horizontal bias of eye movements (Linka et al., 2025). These authors noted that the prolonged development of saccadic anisotropies aligns with our findings on visual field geometry in children (Carrasco et al., 2022) and adolescents –this study– revealing a less pronounced horizontal-vertical asymmetry than adults. We concur with their suggestion that future work should investigate the relation between the development of eye movement patterns and polar angle asymmetries in visual performance.

Adult humans are more sensitive to stimuli presented along the horizontal than the vertical meridian–even within the high-acuity foveola. Interestingly, the VMA is reversed: objects are more easily discerned when presented slightly above than slightly below the center of gaze (Jenks, Carrasco & Poletti, 2025). Future studies should assess whether the HVA and inverted VMA within the foveola are also present in children and chart their developmental trajectories throughout adolescence.

The effects of covert attention and adaptation on polar angle asymmetries have been investigated in adults. Both exogenous (i.e., involuntary; Carrasco et al., 2002; Carrasco et al., 2001; Roberts et al., 2016, 2018; Talgar & Carrasco, 2002) and endogenous (i.e., voluntary; Purokayastha et al., 2021; Tüncok, Carrasco & Winawer, 2025) spatial covert attention, as well as temporal attention (Fernández, Denison & Carrasco, 2019) enhance sensitivity similarly around the visual field, indicating that covert attention does not modulate the extent of the HVA or VMA. In contrast, adaptation has a stronger effect along the horizontal than the vertical meridian, mitigating the HVA in contrast sensitivity and promoting perceptual uniformity around the visual field (Lee & Carrasco, 2025). We are currently investigating how covert attention affects sensitivity in children and adolescents.

A comprehensive account of visual development must consider the interaction between maturation and environmental input. Asymmetries between upper and lower hemifields may relate to functional specialization, where the upper visual field processes distant objects and the lower visual field processes near objects (Previc, 1990). For adults, many tasks requiring manual manipulation involve stimuli in the lower visual field. For children, however, relevant visual information often appears above eye level, and this shifts as they grow.

Nonetheless, these asymmetries are primarily driven by the cardinal meridians rather than the entire upper and lower hemifields. The VMA decreases with polar angle distance from the vertical meridian and is no longer significant at intercardinal locations for contrast sensitivity (Abrams et al., 2012) and acuity (Barbot et al., 2021; reviewed in Himmelberg et al., 2023). Therefore, the specialization for processing distant versus near objects (Previc, 1990) likely contributes to meridional asymmetries but cannot fully account for the polar angle-specific differences observed.

Interestingly, we found that height—after regressing out age—does not correlate with the HVA (**Figure 5A**), but it does correlate with the VMA (**Figure 5B**). These distinct results support the idea that the HVA and VMA are independent and may reflect the spatial statistics of environmental input (Carrasco et al., 2022; Himmelberg et al., 2023; Tünçok, Kiorpes & Carrasco, 2025).

Recently, we discovered that the VMA is inverted in non-human primates: their performance is poorer at the lower, rather than the upper, vertical meridian. Based on this phylogenetic difference, we hypothesize that evolutionary distinctions in locomotion, navigation, and tool use result in differential visuomotor interactions at specific visual field locations (Tünçok et al., 2025).

Characterizing performance around the visual field across development is essential for understanding the functional plasticity of the human brain and how visual systems adapt to their environments. Consistent with evidence that contrast sensitivity (Dekker et al., 2020) and gaze behavior (Linka et al., 2025) continue to mature throughout adolescence, our findings demonstrate that visual field asymmetries in contrast sensitivity also continue to develop, reaching adult-like levels only in the final years of adolescence.

Future research investigating differences in natural scene statistics across age groups—and between humans and non-human primates—could shed light on how environmental factors shape polar angle asymmetries. These findings underscore the importance of conducting behavioral and neurophysiological studies as a function of polar angle to better link anatomical and functional cortical measurements with perceptual performance throughout development. They also highlight the need to account for both age-related and species-specific variations when developing models of vision and brain function.

## RESOURCE AVAILABILITY

### Lead contact

Further information and requests should be directed to the lead contact, Marisa Carrasco.

### Materials availability

This study did not generate new unique reagents.

### Data and code availability

All original experimental code is freely available via an open-access repository (Database: https://github.com/CarrascoLab/PF_adolescents).

Any additional information required to reanalyze the data reported in this paper is available from the lead contact upon request.

## LIMITATIONS OF STUDY

For the present study, we could recruit many more females than males; thus, we lack appropriate statistical power to assess possible biological sex differences. It is an open question if the age effects we report here for the HVA and VMA equally represent both males and females. To minimize the influence of other individual differences, we aimed to obtain large and roughly equal sample sizes for each adolescent age group. Moreover, our participants vary across age and thus some differences contributing to the effect of age could still be related to other individual factors. A longitudinal study assessing both the HVA and VMA would be ideal, but would take many years to complete.

### Author Contributions

MC conceptualized the study, acquired the funding, and supervised and administered the study. CM and MR collected the data. CM analyzed the data and created the figures. MC wrote the original draft. All authors reviewed and edited the manuscript and reviewed the figures.

## Acknowledgments

NIH Grant ODSS/NEI RO1 EY027401 to MC supported this research. We thank the participants in this study, as well as Elizabeth Eberts, Husaifa Fatima, Millie Howard, Imaad Siddiqi, Noa Simoncelli and members of the Carrasco Lab for assistance with subject recruitment and data collection.

## Inclusion and diversity statements

We worked to ensure sex/gender balance and ethnic diversity in the recruitment of human subjects. While citing references scientifically relevant for this work, we also actively worked to promote gender balance in our reference list

## Declaration of generative AI and AI-assisted technologies

None.

## STAR METHODS

## SUBJECT PARTICIPANT DETAILS

Two groups of observers participated in this study: 155 adolescents, 13-17 year-olds (median age = 15.98, mean = 15.64, sd = 1.22; 120 females, 35 males) and 112 adults (median age = 23.79, mean = 28.16, sd = 11.43; 76 females, 32 males, 4 unreported). The data for adults have been previously reported in Carrasco et al. (2022). All participants were recruited from New York City. The adults were undergraduate or graduate students at New York University and other New York City residents. The IRB of New York University approved the study.

## METHOD DETAILS

### Stimuli

Observers viewed a computer display from 57 cm on a 21-in. IBM P260 CRT monitor (1,280 × 960 pixel resolution, 90-Hz refresh rate) calibrated and linearized using a Photo Research (Chatworth, CA, USA) PR-650 SpectraScan Colorimeter. Viewing distance was maintained with a chin-and-forehead rest. On each trial, following a 500 ms fixation period, a target stimulus appeared in one 4 isoeccentric locations centered at 6.2° eccentricity for 80-120 ms (**Figure 1**).

The stimulus was a 4 (84%) or 6 (16%) cycles/degree Gabor patch (a sinusoidal grating embedded in a Gaussian envelope), appearing by itself (35%) or amidst 3 distractors (65%), subtending 2º (19%), 3.2º (65.5%) or 4º (15.5%) of visual angle, oriented 20º (34%) or 30º (66%) from vertical. Previous studies with adults have revealed polar angle asymmetries with all these stimulus parameters (e.g., Abrams et al., 2012; Carrasco et al., 2002; Carrasco et al., 2001, 2002, 2022; Kwak et al., 2024; Purokayastha et al., 2021; Himmelberg et al., 2020). The stimulus contrast was ∼25% for adults and ∼24.5% for adolescents, adjusted individually before and throughout the experiment through an up-down staircase (Levitt, 1971), to yield ∼80% performance accuracy across all locations.

### Procedure

The experiment took place in a darkened and sound attenuated room to provide optimal viewing of the displayed stimuli. Observers viewed the display binocularly. We used a 2-alternative forced-choice orientation discrimination task; performance in this task is contingent upon the observers’ contrast sensitivity and has been used to characterize performance at different locations (e.g., Abrams et al., 2012; Himmelberg et al., 2020; Lee & Carrasco, 2025; Ramirez, Foster & Ling, 2021; Szinte et al., 2018; Tomassini et al., 2015; White et al., 2019).

The experimenter instructed observers to fixate on a central point or cross throughout each trial and to indicate whether the stimulus was tilted right (clockwise) or left (counterclockwise) from vertical with a keypress. A feedback tone indicated whether the response had been correct or incorrect after each trial. Instructions stressed accuracy, and there was no time pressure. Each observer performed 1-2 practice block(s) of 40 trials each, and on average, adults performed 77 trials, and adolescents 47 trials, per location.

Observers’ eyes were tracked to ensure that they maintained fixation throughout the trial using an EyeLink 1000 Desktop Mount eye tracker (SR Research, Ontario, Canada). If observers broke fixation, the trial would be discontinued and replaced at the end of the block, to ensure that stimulus locations and retinal locations were equated.

## QUANTIFICATION AND STATISTICAL ANALYSIS

Discriminability, d’ = z(hit rate) minus z(false-alarm rate), was computed for each observer. A hit was (arbitrarily) defined as a left response to a left orientation and a false alarm as a left response to a right orientation (Jigo & Carrasco, 2020; Hanning, Deubel, & Szinte, 2019; Szinte et al., 2018). The log-linear rule correction was applied while estimating d’ (Brown & White 2005; Hautus, 1995).

Response time (ms) was recorded from stimulus onset until participant’s key press. This secondary dependent variable enabled us to assess any potential speed-accuracy tradeoffs. We calculated the geometric mean for each participant.

In all analyses reported, we collapsed across stimulus parameters, as there was no significant difference in the magnitude of the HVA and VMA for the different stimulus parameters between adults or adolescents. There were no significant interactions for the HVA. When there were interactions for the VMA, it was due to being more pronounced for smaller stimuli (F(2,261) =8.6974, p<0.001) and higher spatial frequencies (F(1,263) =14.141, p<.001).

To evaluate whether height contributes uniquely to the magnitude of the HVA and VMA independently from age, we regressed out age from these measures and then correlated the residuals with height (e.g., Himmelberg et al., 2022; Lee & Carrasco, 2025). We computed semi-partial Spearman correlations by first regressing the HVA and VMA magnitudes on age, extracting the raw residuals, and then correlating those residuals with height.

